# Molecular evolution of terpene synthase underlying the diversification of isoprene emission in Fagaceae

**DOI:** 10.1101/2025.07.31.667835

**Authors:** Yuka Ikezaki, Shuichi N. Kudo, Taichi Nakata, Sora Koita, Ryosuke Munakata, Kazufumi Yazaki, Takeshi Torimaru, Nobuhiro Tomaru, Sachiko Isobe, Hideki Hirakawa, Junko Kusumi, Akiko Satake

## Abstract

Plants emit a wide range of volatile organic compounds, among which isoprene is the most abundant and atmospherically influential. Although oak species are major contributors to isoprene emission, there is considerable variation in isoprene emission capacity within the Fagaceae family. To unravel the evolutionary origins of isoprene emission, we investigated the molecular evolution of terpene synthase (TPS) genes across eight species within the Fagaceae. We identified a Fagaceae-specific TPS-b subclade in which potential isoprene synthase (IspS) activity evolved independently in two gene lineages within subgenus *Quercus*. Ancestral sequence reconstruction revealed that the acquisition of a diagnostic amino acid residue for IspS function arose convergently in these lineages and was subject to positive selection, suggesting adaptive evolution. Ancestral-enzyme assays targeting the gene lineage with high gene expression revealed that the early protein primarily produced monoterpenes from geranyl diphosphate (GPP), whereas their descendants shifted substrate preference to dimethylallyl diphosphate (DMAPP), evolving into dedicated isoprene synthases. Our results indicate that IspS activity was not ancestral in Fagaceae, but evolved approximately 56 million years ago within the subgenus *Quercus*, and has been retained ever since. These findings emphasize the roles of enzyme structural innovation and regulatory shifts in the diversification of volatile terpenoid biosynthesis.

## Introduction

Plants release a wide spectrum of biogenic volatile organic compounds (BVOCs), which include molecules with diverse structural and physicochemical characteristics (Kesselmeier and Staudt, 1999; Laothawornkitkul et al., 2009; Satake et al., 2024). These compounds, emitted from various plant tissues such as leaves, flowers, fruits, and roots, enter the atmosphere in significant quantities (Antonelli et al., 2020). BVOCs serve as a dynamic interface between terrestrial ecosystems and the atmosphere, playing an important role in climate and atmospheric chemistry (Lee Ng et al., 2017). The annual global emission of BVOCs is estimated to be around 1000 Tg C, with terpenoids—including isoprene, monoterpenes, and sesquiterpenes (Guenther, 1995; Guenther et al., 2012). Among these, isoprene alone accounts for roughly half of the total BVOC, underscoring its ecological and atmospheric significance (Guenther, 1995).

In the plant kingdom, isoprene emission is distributed heterogeneously across plant taxa (Dani et al., 2014; Harley et al., n.d.; Monson et al., 2013). It is particularly common in fast-growing plants, especially hardwood trees such as oaks (*Quercus* spp.), *Populus* spp. (Miller et al., 2001), and *Eucalyptus* spp. (Dani et al., 2014). Isoprene emission has also been reported in mosses (Hanson et al., 1999), ferns (Tingey et al., 1987), and legumes (Harley et al., 2004), indicating its occurrence across diverse plant groups. Isoprene is synthesized by isoprene synthase (IspS), a class I terpenoid synthase (TPS) enzyme that converts dimethylallyl diphosphate (DMAPP) into isoprene at a key terminal branching point of the methylerythritol phosphate (MEP) pathway (Sharkey et al., 2013; Silver and Fall, 1995). The TPS family likely originated in ancestral land plants following their divergence from green algae (Jia et al., 2022). In angiosperms, IspS belongs to the TPS-b subfamily (Li et al., 2017; Sharkey et al., 2013). In contrast, recent research has shown that IspS in mosses belongs to the TPS-c subfamily (Kawakami et al., 2023), suggesting that IspS has multiple evolutionary origins within the plant kingdom. In angiosperms, IspS appears to have evolved multiple times, likely from closely related terpene synthases. Among these, *trans*-β-ocimene synthases are considered a key precursor, reflecting the functional diversification of terpene synthases in plants (Li et al., 2017; Sharkey et al., 2013).

An intriguing and well-studied example of the uneven distribution of isoprene emission within a plant clade can be found in the Fagaceae family (Harley et al., n.d.; Monson et al., 2013). In this family, isoprene emission is particularly prominent in the genus *Quercus* (oaks), whereas other genera, such as *Fagus*, *Castanopsis*, and *Lithocarpus* exhibit little to no isoprene emission (Loreto, 2002; Monson et al., 2013; Tani et al., 2024). Intriguingly, within the genus *Quercus*, isoprene emission capacity exhibits significant variation, with a mix of strong isoprene emitters, such as *Q. rubra* and *Q. robur*, and non-emitters, such as *Q. ilex* and *Q. suber* (Loreto and Fineschi, 2015). Based on analyses associating isoprene emission capacity with molecular phylogeny within the Fagaceae, Loreto, 2022 and Monson *et al*., 2013 concluded that isoprene emission was likely an ancestral trait in the genus *Quercus* and was subsequently lost and not restored in certain species within the Quercus sections *Cerris* and *Ilex*. These findings lay a critical foundation for reevaluating the evolutionary history of isoprene emission in the Fagaceae. The growing availability of high-quality genomic and transcriptomic data, even for non-model Fagaceae species (Kremer et al., 2012; Kudo et al., 2025; Satake et al., 2023), presents a valuable opportunity to reconstruct the evolutionary trajectory of isoprene emission by examining genome evolution and the expression patterns of *IspS*.

In the present study, we investigated the evolutionary trajectory of isoprene emission in eight species representing four genera within the Fagaceae family. Using comparative genomics, phylogenetics, and ancestral sequence reconstruction, we uncovered a convergent evolutionary origin of *IspS*, shedding light on a novel evolutionary pathway to isoprene emission in angiosperms.

## Results

### Identification of TPS genes in Fagaceae

To identify TPS genes in Fagaceae, we first obtained genome sequences of eight Fagaceae trees from four genera using public databases (Table S1). These included five species from the genus *Quercus* (*Q. robur*, *Q. lobata*, *Q. rubra*, *Q. acutissima*, and *Q. glauca*), and one species each from the genera *Lithocarpus* (*L. edulis*), *Castanopsis* (*C. hystrix*), and *Fagus* (*F. crenata*) (Table S1). Isoprene emission factors vary significantly among these eight species (Table S2), highlighting their suitability for studying the evolution of isoprene emission. A comparative genomic analysis of these species revealed a highly conserved syntenic structure across genera, including *Quercus*, *Lithocarpus*, and *Castanopsis* (Fig. 1*A*). However, the genomic structure of the genus *Fagus* differed significantly from that of the other genera (Fig. 1*A*), suggesting extensive genome rearrangements following its divergence from the rest of the Fagaceae.

**Figure 1.**
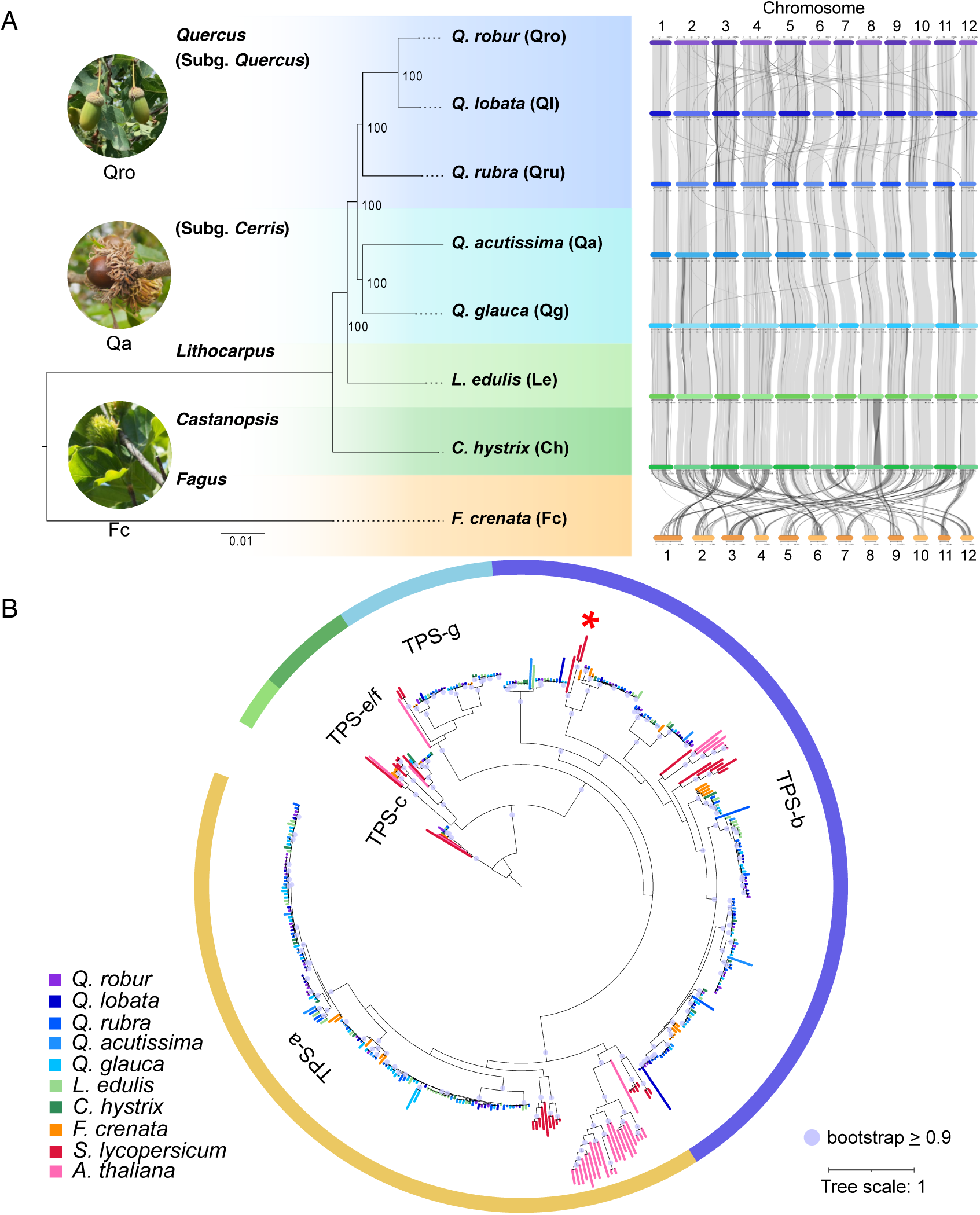
Syntenic relationships of Fagaceae genomes and the phylogenetic analysis of TPS genes. (*A*) Syntenic relationships among eight Fagaceae genomes. The phylogenetic tree was constructed based on whole-genome sequences from eight Fagaceae species, with bootstrap values shown at each branch. Chromosomes of each species are arranged along the corresponding branches of the phylogenetic tree. Syntenic relationships are illustrated with black and gray lines, where black indicates reverse alignments and gray represents standard (collinear) alignments. Chromosome numbers are labeled accordingly. (*B*) Molecular phylogenetic tree of TPS genes in the eight Fagaceae species. TPS genes from *S. lycopersicum* and *A. thaliana* were used as references to define five TPS subclades (TPS-a, -b, -c, -e/f, and -g). The three TPS-b genes in tomato including isoprene synthase were shown by red asterisk. The *ent*-copalyl diphosphate (*ent*-CPP) synthase gene from *Picea sitchensis* was used as the outgroup. The branches whose bootstrap values were 0.9 or more were shown by purple circles.

To identify TPS genes in our target Fagaceae species, we conducted HMMER searches using the annotated protein sequences from the genomes of eight Fagaceae species against the metal-binding domain (PF03936) and the N-terminal domain (PF01397). Each TPS candidate gene was further examined and filtered based on exon-intron structure, including the number and locations of exons (Fig. S1), by comparing with previously reported TPSs (Trapp & Corteau), as well as total sequence length (Fig. S2). The number of identified TPS genes varied across species, ranging from a minimum of 24 in *C. hystrix* to a maximum of 67 in *Q. glauca* (Fig. S3).

The rooted TPS phylogenetic tree constructed from our TPS dataset along with sequences from *Solanum lycopersicum* (tomato) and *Arabidopsis thaliana*, revealed a clear distinction among the TPS-a, TPS-b, TPS-c, TPS-e/f, and TPS-g subfamilies (Fig. 1*B*). TPS-a members in Fagaceae formed a distinct clade, separate from those of tomato and *A. thaliana*, but showing a closer evolutionary relationship to tomato TPS-a genes (Fig. 1*B*). The TPS-c, TPS-g, and TPS-e/f subfamilies were present in all Fagaceae species, except for the TPS-e/f subfamily, which was absent in *Q. acutissima* (Fig. 1*B*).

The TPS-b subfamily represents the largest gene group within Fagaceae species (Fig. 1*B*). The three TPS-b genes in tomato form a distinct clade from other tomato and Arabidopsis TPS-b genes (red asterisk in Fig. 1*B*). These three tomato genes are known to synthesize acyclic monoterpenes, with one of them, TPS47, encoding a cytosolic isoprene synthase (Zhou and Pichersky, 2020).

Many genes in the TPS-a and TPS-b subfamilies are arranged in tandem gene arrays on chromosomes 8 and 9, respectively, suggesting their origin from tandem gene duplication events (Fig. S4*A*). Overall, the genomic locations of these genes were largely conserved across *Quercus*, *Lithocarpus*, and *Castanopsis*. However, in *Fagus*, significant genome rearrangements were observed, indicating a distinct evolutionary trajectory (Fig. S4*B*).

### Isoprene score of TPS-b genes in Fagaceae

To investigate the evolutionary history of *IspS* in Fagaceae, we focused on the TPS-b subfamily that includes known *IspS* from other plant species. We designated the clade containing the tomato *IspS* as TPS-b-target1 (TPS-b-t1) and the clade containing the *Q. serrata IspS* as TPS-b-target1 (TPS-b-t3) (Fig. 2*A*). Between the TPS-b-t1 and TPS-b-t3, we identified a distinct clade, which we designate as TPS-b-target2 (TPS-b-t2) (Fig. 2*A*). Notably, both TPS-b-t2 and TPS-b-t3 consist exclusively of genes from Fagaceae, suggesting that these clades are specific to this family and potentially involved in the evolution of *IspS* in Fagaceae.

**Figure 2.**
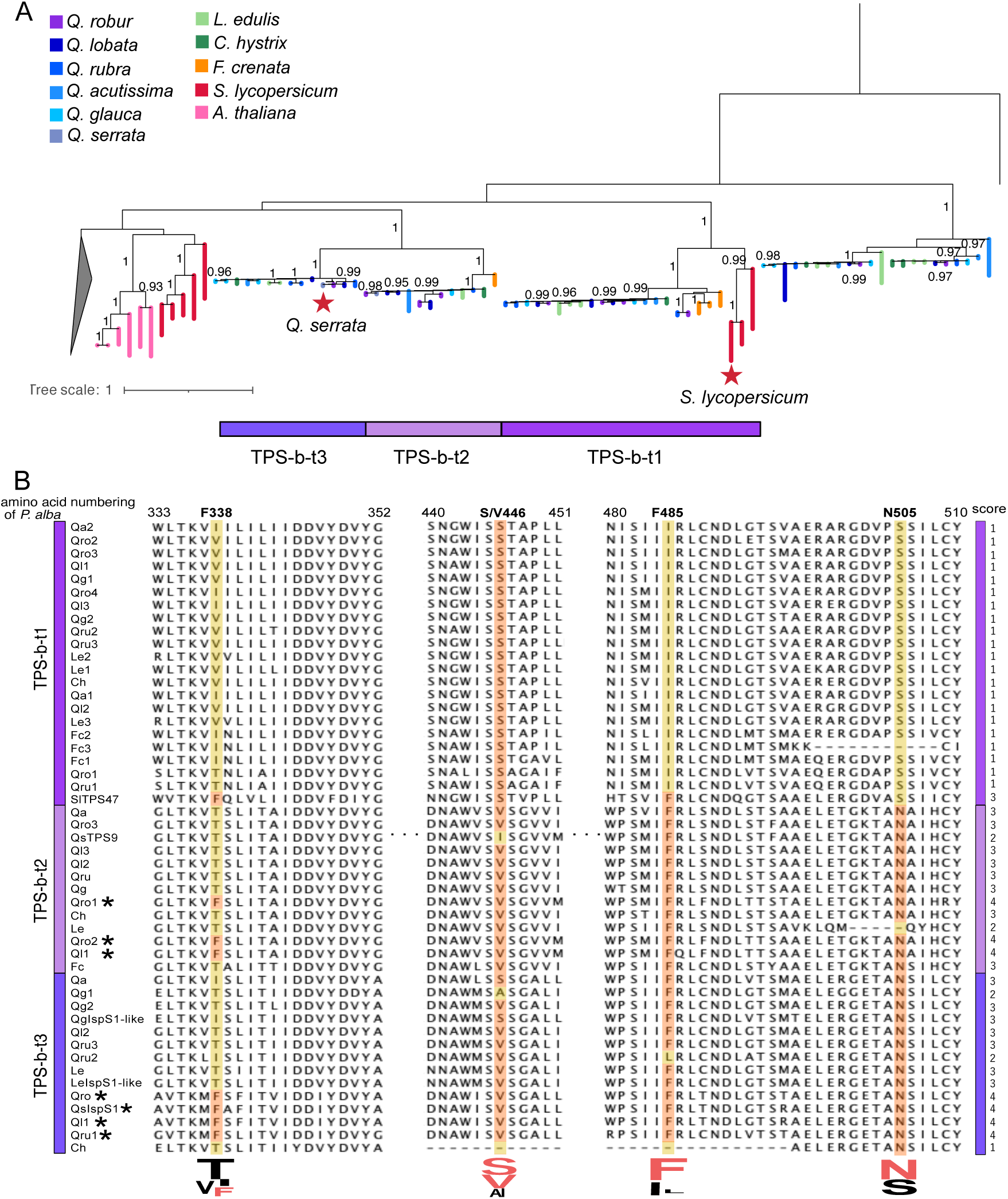
Phylogenetic tree and multiple sequence alignment of TPS genes in the TPS-b Subclade. (*A*) Molecular phylogenetic tree of TPS genes classified within the TPS-b subclade. The phylogeny was inferred with RAxML v8.2.11 under the GAMMA JTT substitution model, using 500 bootstrap replicates to assess node support. A TPS-g gene from *S. lycopersicum* (TPSgSlTPS37) was used as an outgroup. Two genes previously characterized with high isoprene synthase activity—score 3 in *S. lycopersicum* and score 4 in *Q. serrata*—are marked with red stars. (*B*) Multiple sequence alignment of TPS genes from the three TPS-b subclades: TPS-b-t1, TPS-b-t2, and TPS-b-t3. Four conserved amino acid positions used to compute isoprene scores are highlighted by red and yellow shading, red means conserved amino acid at this position and yellow means another. Genes with the highest isoprene score (score 4) are indicated by asterisks.

Sequences of genes included in these three clades were aligned and used to calculate an isoprene score (Fig. 2*B*). The isoprene score serves as a useful screening tool to identify enzymes with exclusive IspS activity based on sequence information and the structural analysis of IspS (Sharkey et al., 2013). This score is calculated using the four conserved amino acid residues (known as the diagnostic tetrad) that are characteristic of IspS. Two of the residues (F338 and F485 in Fig. 2*B*) were suggested to restrict the size of the substrate-binding pocket and play a key role in determining enzyme specificity (Sharkey et al., 2013). While F338 and F485 of the diagnostic tetrad residues are considered highly specific to IspS, the other positions are considered redundant (Li et al., 2017). In IspS in some angiosperm families, valine (V446) is found instead of serine (Li et al., 2017). Therefore, we treated both serine and valine as conserved residues at this position and denoted it as S/V446. The isoprene score ranges from 0 to 4, with a score of 4 representing full conservation of the diagnostic tetrad and thus a high likelihood of exclusive IspS activity.

The multiple sequence alignment revealed that the isoprene score for sequences in the TPS-b-t1 clade was consistently one, whereas the IspS sequence in tomato had a score of 3 (Fig. 2*B*). In the TPS-b-t2 and TPS-b-t3 clades, sequences from the species in the subgenus *Quercus* (*Q. robur*, *Q. lobata*, *Q. rubra*, and *Q. serrata*) were found to have an isoprene score of 4. Specifically, *Q. robur* contained three, *Q. lobata* had two, and *Q. rubra* had one sequence with an isoprene score of 4. In contrast, the maximum isoprene score for sequences from *Lithocarpus*, *Castanopsi*s, and *Fagus* was only three. Many sequences lacked F338, an amino acid considered crucial for isoprene synthase activity (Sharkey et al., 2013). These patterns in isoprene score align with isoprene emission capacities, where *Q. robur*, *Q. lobata*, *Q. rubra*, and *Q. serrata* are known to exhibit high isoprene emission, while other species show lower emission levels (Table S2)

Within the twin clades TPS-b-t2 and TPS-b-t3, *Fagus* sequences were found only in TPS-b-t2, whereas sequences from other species were present in both clades. Most genes within TPS-b-t2 and TPS-b-t3 were located on chromosome 8, with the exception of *F. crenata* (Fig. S4). Interestingly, two genes in *Q. robur* and one in *Q. lobata*, all with an isoprene score of 4, were located on chromosome 3 (Fig. S4), suggesting chromosomal translocation events.

### Seasonal expression patterns of TPS genes

To investigate the seasonal expression patterns of TPS genes, we conducted RNA-seq on leaf and bud tissues collected monthly over a two-year period from *Q. acutissima* growing under natural conditions in Fukuoka, Japan. Additionally, we incorporated previously published field transcriptome data from *Q. glauca* and *L. edulis* (Kudo et al., 2025), which were sampled at the same study site as *Q. acutissima*. On average, 36% and 41% of TPS genes were expressed in leaves (Fig. 3*A*) and buds (Fig. S6*A*), respectively. Within the TPS-b clade, 42% of genes were expressed in leaves, including 6 out of 14 in *Q. acutissima*, 10 out of 28 in *Q. glauca*, and 2 out of 24 in *L. edulis*. Seasonal expression profiles in leaves revealed that most expressed genes exhibited peak expression in summer, although a small fraction of genes showed peaked expression during autumn–winter (e.g., Qa12 and Qg1 in the TPS-a clade in Fig. 3*A*). Notably, expression patterns differed markedly among the three target TPS subclades i.e., TPS-b-t1, t2, and t3. In leaves, only genes in the TPS-b-t3 clade showed clear and consistent summer expression peaks (Fig. 3*D*), whereas expression levels in the TPS-b-t1 and TPS-b-t2 clades were low in all species (Fig. 3*B, C*). These results suggest that TPS-b-t3 genes are primarily responsible for terpenoid biosynthesis. Furthermore, the expression pattern of *Q, serrata IspS* (Koita et al., 2025) within the TPS-b-t3 clade also showed strong expression (Fig. 3*B*-*D*), providing additional support for the functional role of this clade in isoprene biosynthesis. In contrast, in bud tissues, expression levels of genes within the TPS-b-t3 clade were consistently low, and genes in the other two clades, TPS-b-t1 and t2 were relatively high (Fig. S6*B*-*D*), implying a tissue-specific regulation of TPS gene expression.

**Figure 3.**
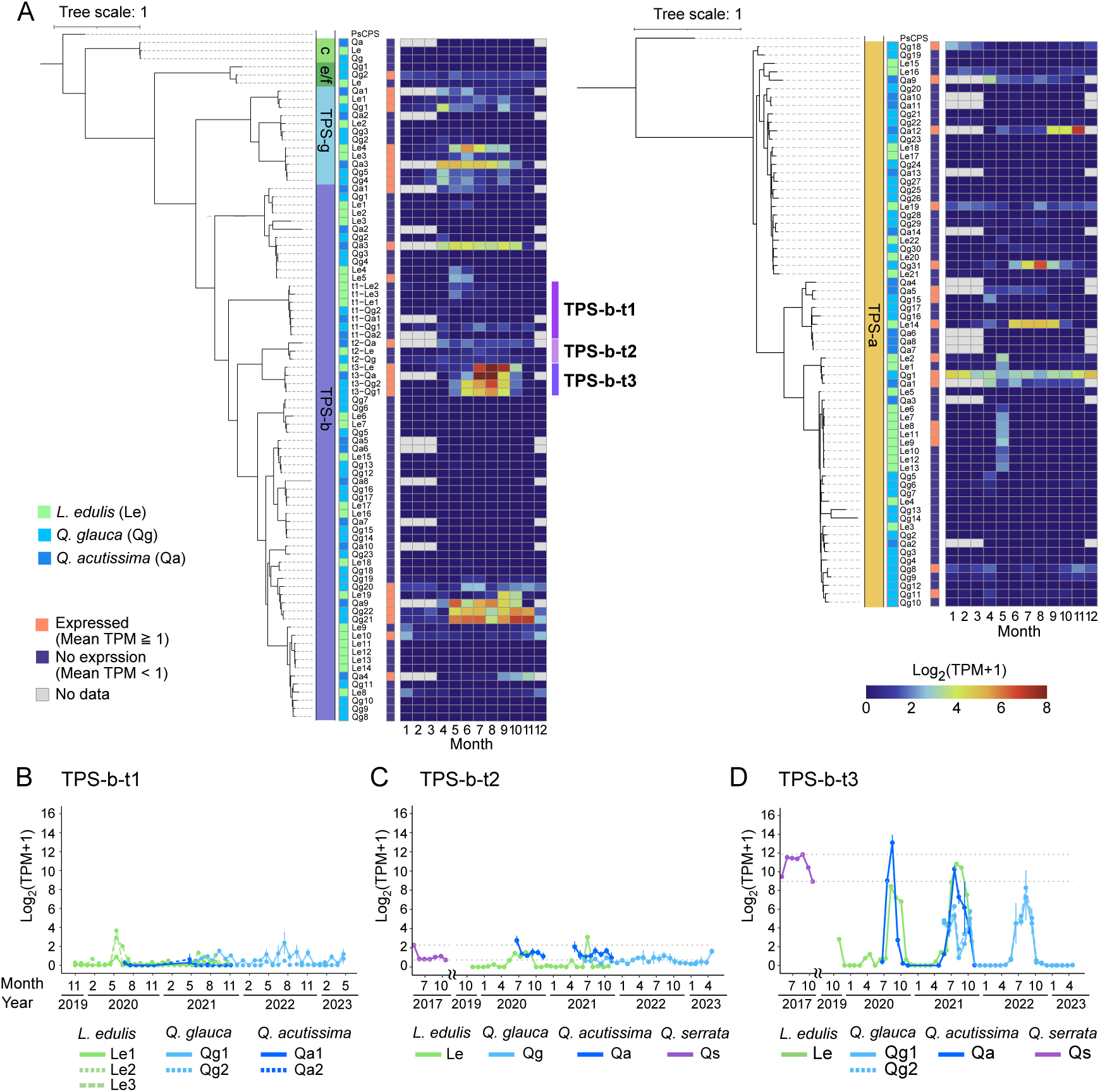
Seasonal expression patterns of TPS genes in leaves. (*A*) Heatmap showing the seasonal expression patterns of TPS genes in three Fagaceae species: *Q. acutissima* (blue), *Q. glauca* (light blue), and *L. edulis* (green). TPS-c genes are not included in the heatmap, as no TPS-c genes were consistently expressed in all three species. Gray squares indicate missing data due to leaf shedding. Genes were considered expressed if the mean of TPM values across all observation periods was ≥1. (*B-D*) Seasonal expression patterns of TPS genes within the subclades TPS-b-t1, TPS-b-t2, and TPS-b-t3. In addition to the three species mentioned above, data from *Q. serrata* were incorporated from a previous study (Koita *et al*., 2025).

### Convergent evolution and positive selection in *IspS* candidates of Fagaceae

To determine whether the genes with an isoprene score of 4 were gained or lost during evolution, we employed the ancestral sequence reconstruction (ASR) method. This approach relies on statistical principles to infer the sequences of putative ancestral genes based on the amino acid or nucleotide sequences of extant species. For this analysis, we used nucleotide sequences from 17 Fagales species belonging to the TPS-b clade (Table S3) and reconstructed putative ancestral sequences using codeml implemented in PAML (https://github.com/abacus-gene/paml).

The estimated ancestral sequences of the last common node of the TPS-b-t1, TPS-b-t2, and TPS-b-t3 clades (node N2; Fig. 4*A*) had an isoprene score of 2, with the conserved residuals S/V446 and F485 (Fig. 4*B*). Subsequently, N505 was gained in the common ancestor of the TPS-b-t2 and TPS-b-t3 clades (node N4; Fig. 4*A*, *B*), prior to their duplication. Following this duplication, F338 was acquired independently in each of the TPS-b-t2 and TPS-b-t3 clades (nodes N6 and 8; Fig. 4*A B*), specifically in the lineage leading to the three species within subgenus *Quercus* (*Q. robur*, *Q lobata*, and *Q. rubra*).

**Figure 4.**
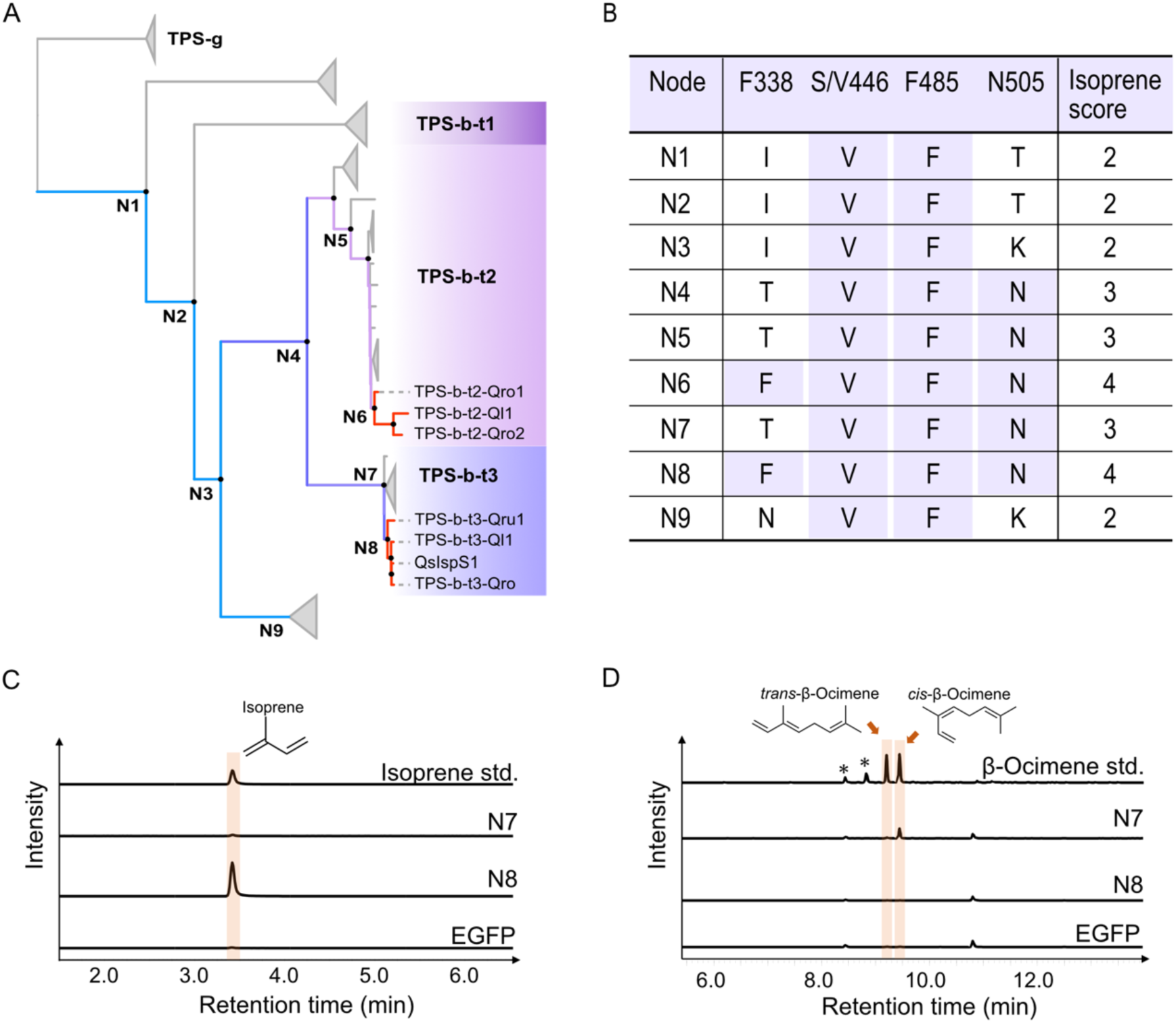
Results of ancestral sequence reconstruction of Fagaceae TPS genes using PAML. (*A*) Maximum-likelihood phylogeny of the TPS gene set used for ancestral reconstructions. The phylogeny was inferred with RAxML v8.2.11 under the GAMMA GTR substitution model, using 1,000 bootstrap replicates to assess node support. Internal nodes N1–N8 mark the bifurcation points analysed. Branch colours denote the inferred isoprene score—blue (2), purple (3) and red (4). The three focal clades (TPS-b-t1, TPS-b-t2 and TPS-b-t3) are highlighted in distinct colours for ease of reference. (*B*) Inferred amino-acid states at the diagnostic tetrad and the corresponding isoprene score for each ancestral node (N1–N8). The posterior probability (PP) of the most likely codon/amino-acid state was as follows: N1 – S/V446: GCA (Ala, PP = 0.263) → GTA (Val, PP = 0.600); F485: CTT (Leu, PP = 0.523) → TTT (Phe, PP = 0.405). N4 – T505: AAA (Lys, PP = 0.774) → AAT (Asn, PP = 0.956). N6 – F338: ACT (Thr, PP = 0.999) → TTT (Phe, PP = 0.970). N8 – F338: ACT (Thr, PP = 0.996) → TTT (Phe, PP = 0.990). (*C*) and (*D*) Enzyme assays of the ancestral sequence products using DMAPP and GPP as substrates. The reaction products were detected at *m/z* 67 for isoprene (*C*) and *m/z* 93 for monoterpenes (*D*) as the substrate. The *cis* or *trans* configurations of β-ocimene were identified based on spectral annotation. The chromatograms, except for the β-ocimene standard, are plotted on the same scale. Asterisks represent peaks of impurities.

Notably, the gain of F338 occurred independently in both the TPS-b-t2 and the TPS-b-t3 clades, despite requiring two nucleotide substitutions for the transition from ACT (T) to TTT (F). We also employed GRaSP (https://github.com/charles-abreu/GRaSP) to infer the ancestral sequences and reached the same conclusion that the gain of F338 occurred independently in both the TPS-b-t2 and TPS-b-t3 clades (Fig. S7). Although the inferred ancestral sequences for the distant ancestor at nodes N1–N3 differed slightly between PAML and GRaSP, the evolutionary pattern across the remaining nodes was consistent (Fig. S7). These results suggest that the acquisition of isoprene-producing function occurred through the stepwise accumulation of mutations, potentially driven by convergent molecular evolution in the TPS-b-t2 and TPS-b-t3 clades.

We next tested for evidence of positive selection in the clades that acquired F338 (predefined as foreground branches) by applying the branch-site model in codeml (Yang, 2007; Zhang et al., 2005). This approach identifies amino acid sites where the nonsynonymous to synonymous substitution rate ratio (ω) exceeds one, indicative of positive selection. The analysis revealed significant evidence of positive selection in the foreground branches (likelihood ratio test, *P* = 1.027×10⁻⁶) with four residuals identified as candidates under positive selection (Bayes Empirical Bayes posterior probability > 0.95) (Table S4; Fig. S8). Notably, the residue corresponding to F338 was included among these candidates, suggesting that the convergent gain of this residue in the TPS-b-t2 and TPS-b-t3 clades was likely driven by positive selection.

### *in vitro* enzyme assays demonstrated that the *IspS* evolved from monoterpene synthase

To evaluate the evolutionary trajectory of enzymatic function, ancestral TPS sequences were artificially synthesized (Dataset S1) and subjected to enzymatic characterization using an *Escherichia coli* expression system. Because genes within the TPS-b-t3 clade exhibited high expression levels, we focused our analyses on this clade. Enzyme assays using purified recombinant proteins were conducted with either DMAPP (for IspS activity) or GPP (for mono-TPS activity) as a substrate. The ancestral enzyme at node N7 exhibited mono-TPS activity, specifically producing *trans*- and *cis*-β-ocimene, while the enzyme at node N8 exhibited IspS activity only (Fig. 4*C*, *D*). Ancestral proteins reconstructed using both PAML and GRaSP showed identical enzymatic activities at nodes N7 and N8 (Fig. 4*C*, *D*; Fig. S7*C*, *D*). These results indicate that *IspS* with an isoprene score of 4 evolved from β-ocimene synthases, supporting the functional shift during evolution.

## Discussion

Using comparative genomics, ancestral-sequence reconstruction, and *in vitro* assays of ancestral enzymes, we found that *IspS* candidates with an isoprene score of 4 arose independently twice in the subgenus *Quercus*—once in the TPS-b-t3 clade and once in the TPS-b-t2 clade—within the TPS-b subfamily in Fagaceae. Because the subgenus *Quercus* diverged from its common ancestor roughly 56 million years ago (Zhou et al., 2022), we infer that the onset of substantial isoprene emission in Fagaceae to the Eocene. Moreover, in all species we examined (*Q. robur*, *Q lobata*, and *Q. rubra*), once the IspS activity evolved, it has been retained. This contrasts with rosids, where a single early gain of isoprene synthase was followed by multiple independent losses (Shang et al., 2024). Fagaceae therefore provide a complementary pattern—multiple, convergent gains coupled with long-term retention—highlighting diverse evolutionary routes to isoprene emission in angiosperms.

The early TPS-b-t3 proteins preferred GPP and mainly produced β-ocimene, whereas their descendants evolved to favour DMAPP and became dedicated isoprene synthases in the subgenus *Quercus*. The shift would correlate with pocket-narrowing substitutions—most notably the substitution from Thr to Phe at the 338 residue—that exclude the larger GPP substrate. These results confirmed the previous view in which the ocimene synthases is ancestral TPS for which IspS evolved by active site size reduction following acquisition of F338 (Gray et al., 2011). The combination of branch-level and site-level likelihood tests detected positive selection at this residue, indicating adaptive convergence toward isoprene emission. We also identified three additional residues under positive selection, suggesting that other sites may contribute to isoprene emission, although the effects of these sites on substrate specificity are currently unknown. Isoprene provides a threefold heat shield—membrane stabilization, ROS scavenging, and Ca²⁺-mediated heat-shock signaling (Zuo et al., 2025)—mechanisms that likely protect photosynthesis and sustain growth in *Quercus* species with thin, deciduous leaves under rising temperatures.

Terpenoids represent the largest and most diverse class of natural products in terrestrial plants (Chen et al., 2011). Comparative and functional genomics can chart the continuous evolutionary innovation of terpenoid-biosynthetic pathways with unprecedented resolution. Such insight will deepen our understanding of how chemical diversity has shaped plant adaptation and diversification on the earth.

## Materials and Methods

### Genome synteny analysis

To compare the genome structures of Fagaceae species across genera, published genome sequences of eight Fagaceae species from four genera were obtained from the public databases and used for subsequent analyses (Table S1). We first performed an all-vs-all BLASTp search using protein sequences with a cut-off e-value of 1E-5 to identify homologous relationships across the genomes. To investigate genomic synteny, we detected syntenic blocks within or between genomes using MCScanX (https://github.com/wyp1125/MCScanX), based on the results of the above BLASTp searches. The synteny blocks were visualized using SynVisio (https://synvisio.github.io). To clarify syntenic relationships across chromosomes in each species, we renumbered chromosomes according to the chromosomal order in *Q*. *robur*. Additionally, chromosomes oriented opposite to those of *Q. robur* were reversed. These adjusted chromosomes were used for subsequent analyses.

### Phylogenetic analysis of TPS genes

Alignment and phylogenetic tree construction were conducted using the build function of ETE3 v3.1.3 (https://etetoolkit.org). Sequence alignment was performed with MAFFT v6.861b (https://mafft.cbrc.jp), alignment trimming with trimAl v1.4.rev (https://vicfero.github.io/trimal), and phylogenetic tree construction with RAxML v8.2.11 (https://cme.h-its.org/exelixis/web/software/raxml) using the GAMMA JTT model and 500 bootstrap replications. The resulting phylogenetic tree was visualized using the online tool iTOL (https://itol.embl.de). The analyzed TPS genes were classified into five subclade groups: TPS-a, TPS-b, TPS-c, TPS-e/f, and TPS-g, based on reference genes from Arabidopsis and tomato. The chromosomal locations of these classified genes were visualized using TBtools-II v2.2.10 (https://github.com/CJ-Chen/TBtools-II).

### Identification of TPS family members

We retrieved candidate genes of the TPS family based on two conserved motifs: the metal-binding domain (PF03936) and the N-terminal domain (PF01397). Protein sequences from eight Fagaceae species were used for a domain search against the Pfam-A database ver 35.0 (Mistry et al., 2021) using HMMER ver 3.3.2 (http://hmmer.org) with a cut-off e-value of 1E-5. For genes with more than two isoforms, the longest isoform was selected.

To exclude candidate genes with short sequence lengths, we utilized the conserved exon-intron arrangement patterns found in TPS genes (Trapp and Croteau, 2001). Across a broad range of plant species, these patterns are well-documented and categorized into three classes: Class I (12–14 introns), Class II (9 introns), and Class III (6 introns) (Trapp & Croteau, 2001). Notably, Class II TPS genes are specific to gymnosperms, meaning that only Class I and Class III TPS genes are present in the angiosperm species analyzed in this study. To identify putative TPS genes, we generated histograms depicting the intron count for each gene within each species. The majority of genes exhibited exon-intron patterns consistent with either Class I or Class III (Fig. S1). Based on this distribution, we selected genes containing more than five exons.

Furthermore, some genes remained too short for detailed analysis. To address this, genes encoding proteins shorter than 400 amino acids were excluded (Fig. S2). To classify the subgroups of TPS genes, we incorporated protein sequences of well-characterized TPS genes from *S*. *lycopersicum* and *A*. *thaliana* as references for phylogenetic analysis. These reference sequences were obtained from Table S7 in Zhou and Pichersky (Zhou and Pichersky, 2020).

Additionally, *IspS* and *IspS*-*like* from *Q*. *serrata* were obtained from previous study (Koita et al., 2025)and included in the analysis. To root the phylogenetic tree, we included the *ent-copalyl diphosphate synthase* sequence from *Picea sitchensis* as an outgroup.

### Phylogenetic analysis of TPS genes

Alignment and phylogenetic tree construction were conducted using the build function of ETE3 v3.1.3 (https://etetoolkit.org). Sequence alignment was performed with MAFFT v6.861b (https://mafft.cbrc.jp), alignment trimming with trimAl v1.4.rev (https://vicfero.github.io/trimal), and phylogenetic tree construction with RAxML v8.2.11 (https://cme.h-its.org/exelixis/web/software/raxml) using the GAMMA JTT model and 500 bootstrap replications. The resulting phylogenetic tree was visualized using the online tool iTOL (https://itol.embl.de). The analyzed TPS genes were classified into five subclade groups: TPS-a, TPS-b, TPS-c, TPS-e/f, and TPS-g, based on reference genes from Arabidopsis and tomato. The chromosomal locations of these classified genes were visualized using TBtools-II v2.2.10 (https://github.com/CJ-Chen/TBtools-II).

### Isoprene synthase gene analysis

To investigate molecular evolution of *IspS* in Fagaceae, we utilized published *IspS* sequences from *S*. *lycopersicum* (Zhou and Pichersky, 2020) and *Q. serrata* (Koita et al., 2025), both belonging to the TPS-b subfamily. Since an adjacent subclade was located near the subclades containing these sequences, we focused on the three subclades that include 44 Fagaceae genes.

These genes of three subclades were aligned by MAFFT v7.520 (https://mafft.cbrc.jp) with the reference *IspS* sequence from *Populus alba* (AB198180) (Sasaki et al., 2005), *S. lycopersicum* (Zhou and Pichersky, 2020)and *Q. serrata* (Koita et al., 2025) to calculate an isoprene score (Sharkey et al., 2013).

### Sample collection for RNA sequencing

To evaluate expression of TPS genes in natural conditions, we analyzed transcriptome data obtained from leaf and bud tissues in three Fagaceae species, *Q. acutissima*, *Q. glauca*, and *L. edulis*. We generated data for *Q. acutissima*, whereas those for *Q. glauca* and *L. edulis* were obtained from the previous study (Kudo et al., 2025). For all data, samples were collected every four weeks spanning from 2020–2021 in *Q. acutissima*, 2022–2023 in *Q. glauca* and 2019–2021 in *L. edulis* in Fukuoka prefectures in Japan. For each species, we targeted three individuals and each sample was collected from three branches per individual, except for *L. edulis*. Given the consistent expression patterns observed among individuals in a previous study (Satake et al., 2023), we focused on only one individual for *L. edulis*. Study sites for *Q. acutissima* and *Q. glauca* is the biodiversity reserve on the Ito campus of Kyushu University (33°35′ N, 130°12′ E, 20 to 57 m a.s.l). *L. edulis* were studied at the Imajyuku Field Activity Center (33°33′ N, 130°16′ E, 84 to 111 m a.s.l). Leaf samples were collected from spring to summer in the deciduous species *Q. acutissima*, whereas in the evergreen species *Q. glauca* and *L. edulis*, sampling was conducted throughout the year. RNA-seq samples were collected and preserved in 2 ml microtubes containing 1.5 ml of RNA-stabilizing reagent (RNAlater; Thermo Fisher Scientific, Waltham, MA, USA), then stored at −80°C until RNA extraction, following the protocol described by Kudo *et al*., 2025.

### RNA extraction and RNA-seq analysis

Total RNA was extracted from leaf samples following the protocol described previously (Miyazaki et al., 2014). RNA integrity was assessed using the method outlined previously (Kudo et al., 2025). For each sample, 5–6 µg of total RNA was sent to Hangzhou Veritas Genetics Medical Institute Co., Ltd., where cDNA libraries were prepared using the NEBNext Ultra II RNA Library Prep Kit for Illumina. Transcriptome sequencing (150 bp paired-end reads) was performed on an Illumina NovaSeq 6000 platform (Illumina, San Diego, CA, USA). Gene expression quantification followed a three-step pipeline: (1) quality filtering was performed using fastp v0.20.0 with default settings (https://github.com/OpenGene/fastp); (2) RNA-seq reads were aligned to the reference genome using TopHat v2.1.1 (https://ccb.jhu.edu/software/tophat/index.shtml); and (3) transcript abundance was quantified using StringTie v1.3.5 (https://ccb.jhu.edu/software/stringtie).

### Seasonal expression patterns of TPS genes in Fagaceae

To investigate the seasonal expression patterns of TPS genes, we extracted TPS gene transcripts from RNA-seq data of *Q. acutissima*, *Q. glauca*, and *L. edulis*. For each gene, we calculated the mean expression level in transcripts per million (TPM) across three individuals, except for *L. edulis*, for which data were available from only one individual (Kudo et al., 2025). TPM values were transformed using log₂(TPM + 1). Genes with a mean TPM > 1 across all samples were considered to be expressed. A heatmap was generated in R ver. 4.3.1 using the ‘pheatmap’ package, with gene order arranged according to TPS clades (Fig. 3*A*). In addition to these three species, we also obtained expression data for *IspS* and *IspS*-like in *Q. serrata* (Koita et al., 2025), which were used to plot seasonal changes in expression levels. To generate the heatmap, we calculated the average gene expression levels for each month by aggregating data from different individuals over a two-year period for each species.

### Ancestral amino acid sequence inference

To infer the ancestral amino acid sequences of putative *IspS* in *Quercus* species, we analyzed 95 representative nucleotide sequences, which have no ambiguity, of TPS-b subfamily members from Fagales. These sequences include genes identified through the genomic survey of eight Fagaceae species described above, genes from *Q. serrata*, and additional genes retrieved via BLAST searches against the database. The BLAST search identified genes from *Q. suber*, *L. litseifolius*, *Castanea mollissima*, *Juglans regia*, *J. maicrocarpa x J. regida*, *Carya illinoinensis*, *Corylus avellana*, and *Alnus glutinosa*, resulting in a dataset comprising genes from 17 Fagales species. (Table S3). Additionally, seven TPS-g subfamily genes were used as an outgroup.

Sequence alignment was performed using MACSE v2 (https://www.agap-ge2pop.org/macse), which can account for the underlying codon structure of protein-coding sequences and generate both nucleotide (NT) and amino acid (AA) alignments. Phylogenetic reconstruction and ancestral amino acid sequence inference were conducted separately for NT and AA alignments. ML phylogenetic trees were constructed with RAxML v8.2.11 (https://cme.h-its.org/exelixis/web/software/raxml) with 1000 bootstrap replications. The GAMMA GTR model (GTR+I+G) and the GAMMA JTT model (JTT+I+G) were used for the NT and AA alignment, respectively. The resulting ML trees exhibited compatible topology, particularly the splits of the major clusters. Two strongly supported clusters of *IspS* with the maximum score were identified in both trees. Ancestral amino acid sequences were inferred at each internal node using codeml in PAML (https://github.com/abacus-gene/paml) with the F3x4 codon model for the NT alignment. We also employed GRaSP (https://github.com/charles-abreu/GRaSP) to infer the ancestral sequences from the AA alignment and estimate the location of gaps in the ancestral sequences.

### Detection of positive selection

To investigate whether amino acid sites in putative *IspS* (foreground) have undergone positive selection compared with the other TPS-b subfamily members (background), we applied the branch-site test for positive selection (Zhang et al., 2005). The NT alignment and MLtree used for the ancestral sequence inference were employed in this analysis. Alignment gaps were removed prior to the analyses. Previous studies have shown that gene conversion and recombination events could lead to false positives (Anisimova et al., 2003; Casola and Hahn, 2009). To account for this, we applied the GARD method implemented in HYPHY (https://hyphy.org) to detect potential gene conversion or recombination events. No significant evidence of gene conversion or recombination was found in the genes, so subsequent analyses were conducted using the full-length alignments.

All branches within two clades of the genes with the isoprene score of 4 were set as foreground branches and the branch-site tests were conducted using codeml (Yang, 2007). The null model was set to model = 2, NSsites = 2, fix_omega = 1 and omega = 1, and the alternative model (Model A) was with model = 2, NSsites = 2 and fix_omega = 0 (to estimate ω₂). The Bayes empirical Bayes (BEB) method was used to identify sites potentially under positive selection on the foreground branches.

### Materials and reagents for *in vivo* enzymatic characterization

DMAPP and GPP were used as substrates, with DMAPP purchased from TargetMol (Boston, USA) and GPP from Sigma-Aldrich (St. Louis, USA). As standard compounds, isoprene was obtained from Tokyo Chemical Industry Co., Ltd. (Tokyo, Japan), and β-ocimene was purchased from Cayman Chemical (Ann Arbor, USA).

### Construction of *E. coli* expression vectors

To express the ancestral sequences estimated at nodes N7 and N8 by PAML (*N7_paml* and *N7_paml*) and GRaSP (*N7_grasp* and *N8_grasp*) in *E. coli*, their putative transit peptides at N-terminus were removed and the truncated CDSs were further codon-optimized for *E. coli*. The optimized sequences were synthesized by GeneArt (Thermo Fisher Scientific). These sequences were amplified by PCR using KOD One® (Toyobo, Osaka, Japan) with the primer pairs shown in Table S5. The amplification program is composed of an initial denaturation at 94°C for 1 min followed by 40 cycles of denaturation at 98°C for 10 sec, annealing at 51°C for 5 sec, and extension at 68°C for 10 sec, with a final extension at 68°C for 5 min. The amplicons were individually inserted into pET22b(+) vector (Novagen, Madison, US) by In-Fusion reaction (Takara Bio, Kusatsu, Japan) using NdeI and BamHI sites, which enables the production of proteins with the histidine tag (6×His). The sequence of *N8_paml* was constructed by introducing a single mutation to *N8_grasp* by PCR using KOD One® (Toyobo) and the primer pair *N8_paml*_Fw and *N8_paml*_Rv. This amplification was carried out with an initial denaturation at 94°C for 1 min, followed by 40 cycles of denaturation at 98°C for 10 sec, annealing at 51°C for 5 sec, and extension at 68°C for 40 sec, with a final extension at 68°C for 5 min. The amplicon was introduced into the pET22b(+) vector in the same process as described above.

### Enzymatic characterization

Constructs *ΔTP-N7_paml-6×His*, *ΔTP-N7_grasp-6×His*, *ΔTP-N8_paml-6×His*, and *ΔTP-N8_grasp-6×His* were individually introduced into *E. coli* BL21 (DE3) for recombinant protein production. Protein preparation was performed as previously described (Koita *et al*., 2025). Protein concentrations were measured using a Qubit® 2.0 Fluorometer and the Qubit Protein Assay Kit (Thermo Fisher Scientific).

A standard reaction mixture (200 µl) contained 0.5 mM DMAPP or GPP, 20 mM MgCl₂, and 160 µl of purified enzyme. For assays using DMAPP, 27–32 μg of enzyme were used, whereas 1.4–1.7 μg were used for GPP-based assays. Reactions were incubated at 37 °C for 30 minutes in 2 ml glass vials (Thermo Fisher Scientific). GC-MS analysis of the enzymatic products was carried out as described in Koita *et al*., 2025.

## Acknowledgments

We thank Kayoko Ohta for her technical support with RNA extraction. We also thank Koharu Yamaguchi, Ryota Aoyagi, and Ryosuke Imai for their help with sample collection for RNAseq, Sangwan Kim for his support with registration of RNA sequence data to DDBJ, and Kenji Fukushima for his insightful comments on the bioinformatic analyses.

## Additional information

## Data availability

The RNAseq data that support the findings of this study are openly available in DDBJ, Bioproject ID: PRJDB35868.

## Author Contributions

Y.I. and A.S. designed research; Y.I., S.N.K., N.T., S.K., R.M., K.Y., S.I., H.H., J.K. and A.S. performed research; Y.I., S.N.K., N.T., S.K., R.M., K.Y., S.I., H.H., J.K. and A.S. analyzed data; T.T. and N.T. provided genome data; Y.I. and A.S. wrote the paper with inputs from all the authors.

## Additional files

**Supplementary Information.** Supplementary figures (Fig. S1 to Fig. S8) and supplementary tables (Table S1 to S5) were included int this file.

Supplementary 1.

**Supplementary Dataset.** Supplementary dataset (Dataset S1) was included in this file. It is multiple sequence alignment of ancestral sequences of Fagaceae TPS genes using PAML and GRaSP.

Dataset S1.

