## Supplementary 1 for "Molecular evolution of terpene synthase underlying the diversification of isoprene emission in Fagaceae"

The following Supporting Information is available for this article:

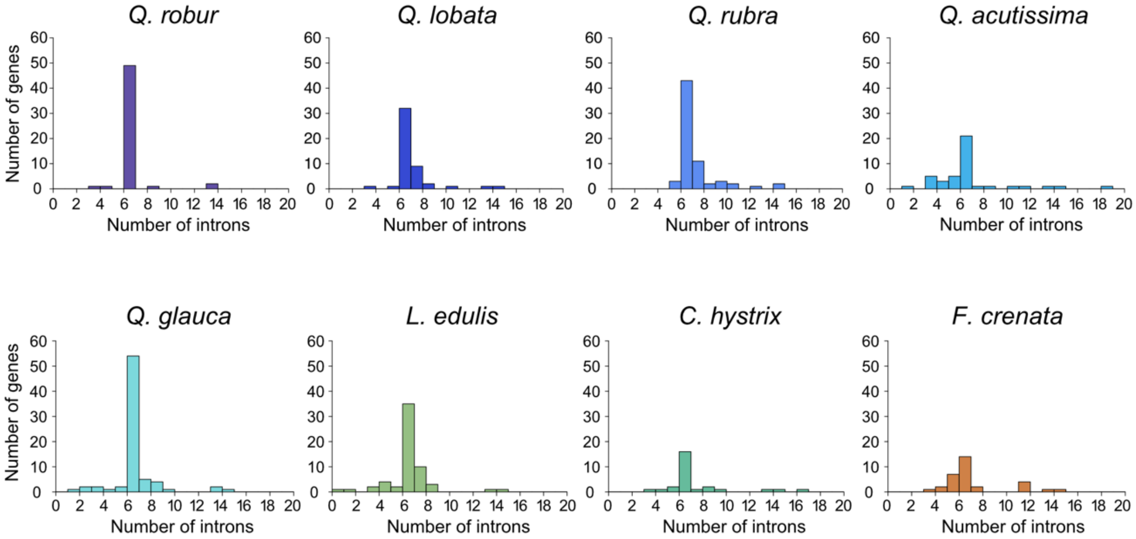


**Figure S1.** Distribution of intron numbers in TPS candidate genes across eight Fagaceae species.

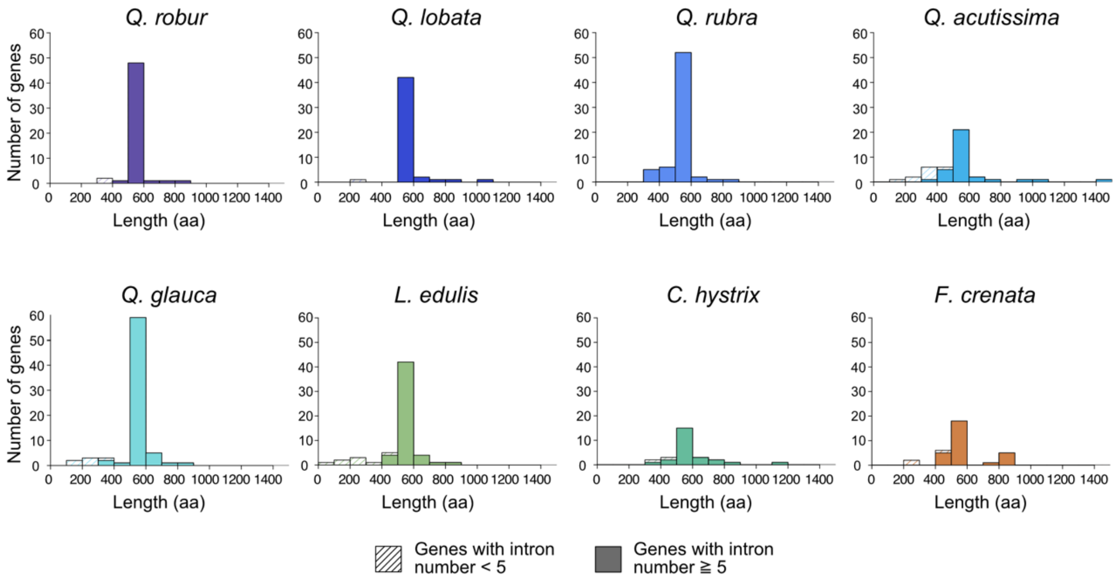


**Figure S2.** Histogram of amino acid lengths of TPS genes in eight Fagaceae species. Cells with diagonal lines represent all TPS genes, while filled cells indicate those retained after filtering out genes with low intron numbers (fewer than five). Genes shorter than 400 amino acids were excluded from subsequent analyses.

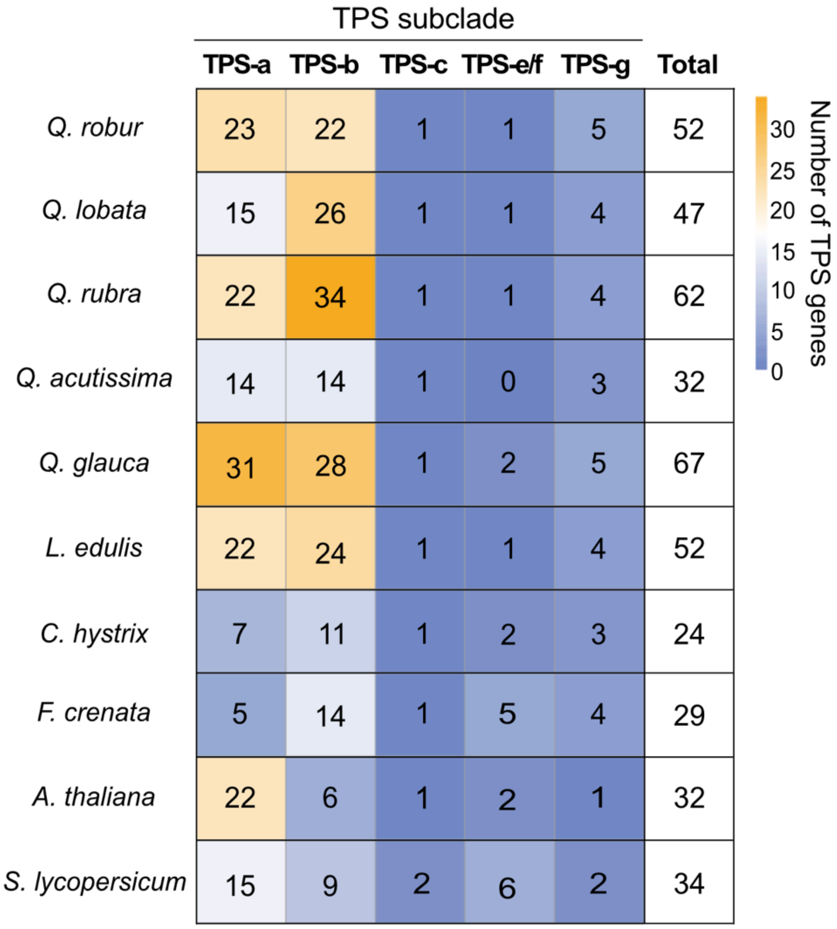


**Figure S3.** Number of TPS genes in each subclade of Fagaceae genomes. The heatmap shows the number of TPS genes assigned to each category, with warmer colors representing higher gene counts. TPS gene counts from *Arabidopsis thaliana* and *Solanum lycopersicum* are also included for comparison.

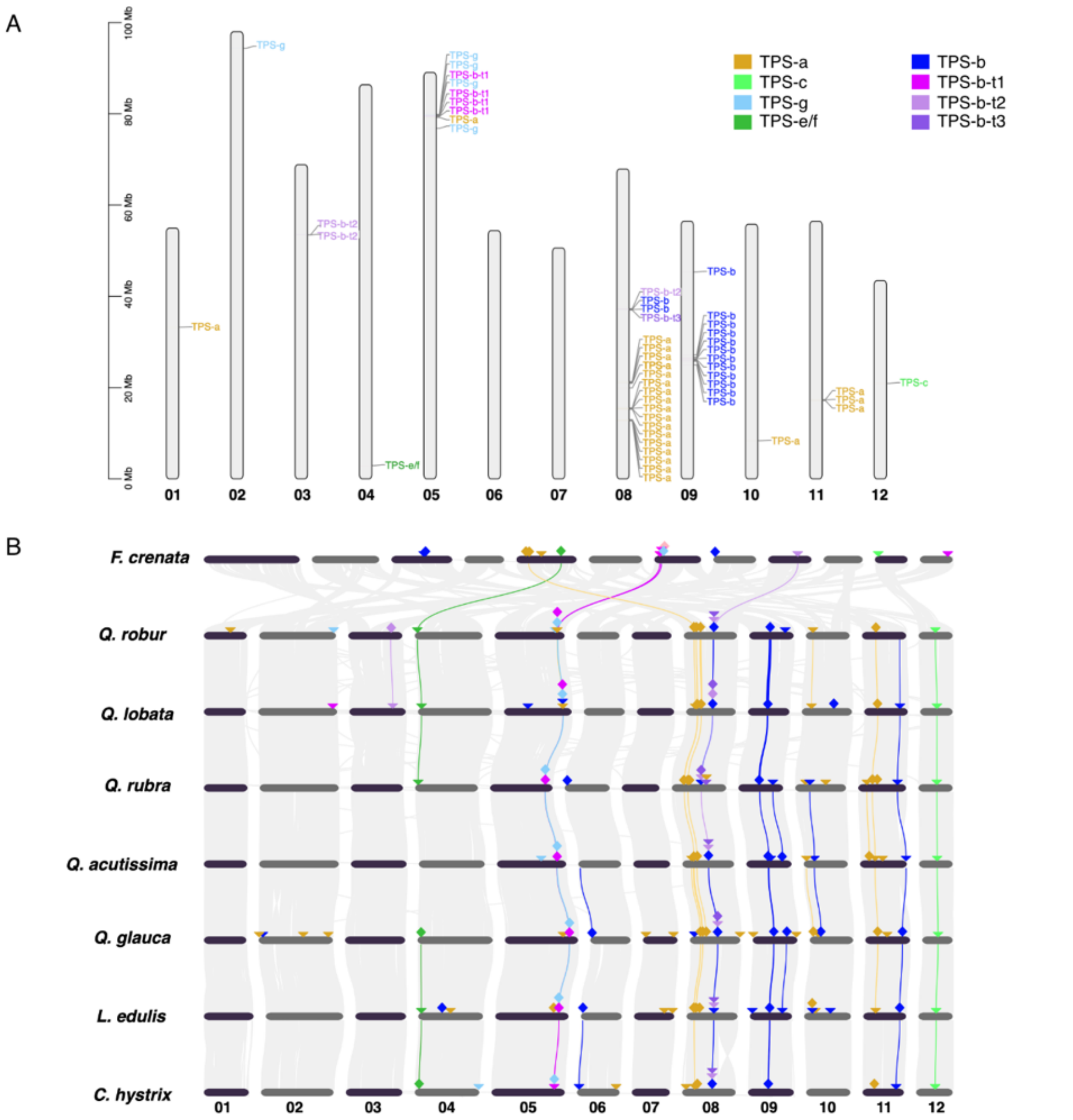


**Figure S4.** Genomic distribution of TPS genes in eight Fagaceae species. (*A*) Chromosomal distribution of TPS genes in the twelve chromosomes of *Quercus robur*. (*B*) Collinearity plot of TPS genes across the twelve chromosomes of eight Fagaceae species. Colored inverted triangles indicate the locations of TPS genes classified into different subclades. Diamonds mark regions where multiple TPS genes (two or more) are clustered. Colored lines represent collinear genomic blocks between species that include TPS genes, highlighting conserved syntenic relationships.

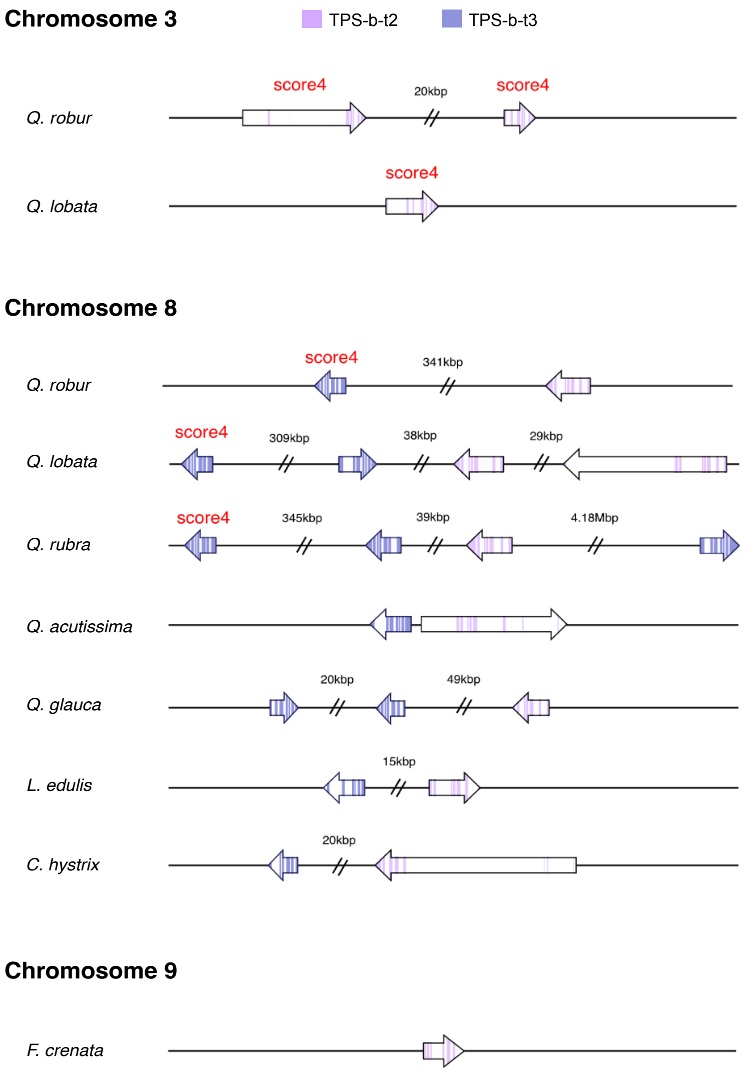


**Figure S5.** Chromosomal locations of TPS-b-t2 and TPS-b-t3 subclade genes in Fagaceae species. In seven species (excluding *Fagus crenata*), all TPS-b-t2 genes were located on chromosome 8. In *Quercus robur* and *Q. lobata*, TPS-b-t2 genes were found on both chromosomes 8 and 3. In contrast, TPS-b-t2 genes in *F. crenata* were located on chromosome 9. A collinearity block linking the distal region of chromosome 8 in *Q. robur* (which harbors TPS-b-t2 genes) and chromosome 9 in *F. crenata* is shown in Fig. S3*B*, indicating a conserved genomic region. Colored segments of arrows represent gene exons.

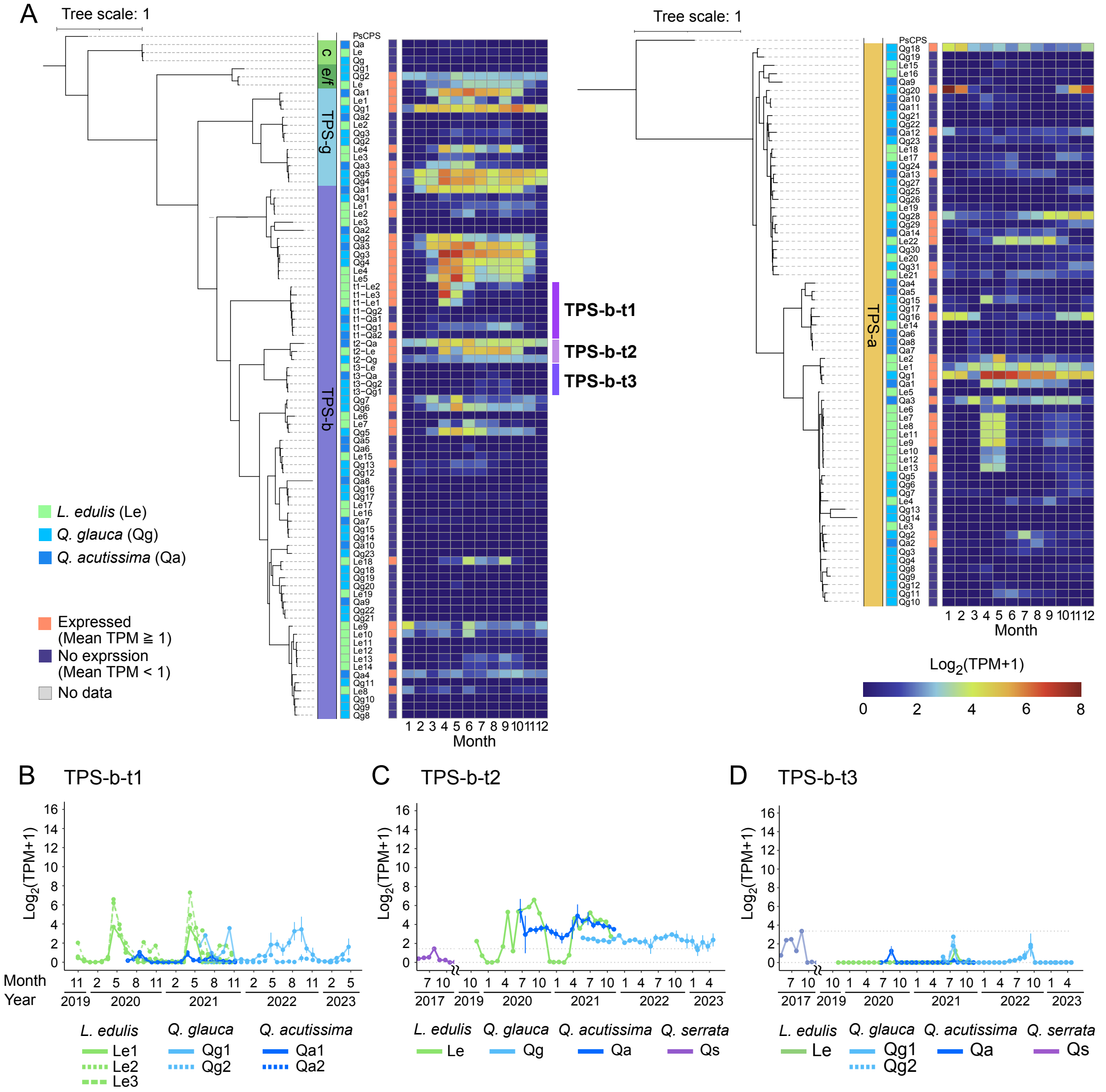


**Figure S6.** Seasonal expression patterns of TPS genes in buds. (*A*) Heatmap showing the seasonal expression patterns of TPS genes in three Fagaceae species: *Quercus acutissima* (blue), *Q. glauca* (light blue), and *Lithocarpus edulis* (green). TPS-c genes are not included in the heatmap, as no TPS-c genes were consistently expressed in all three species. Genes were considered expressed if the mean of TPM values across all observation periods was ≥1. (*B-D*) Seasonal expression patterns of TPS genes within the subclades TPS-b-t1, TPS-b-t2, and TPS-b-t3. In addition to the three species mentioned above, data from *Q. serrata* were incorporated from a previous study (Koita et al., 2025).

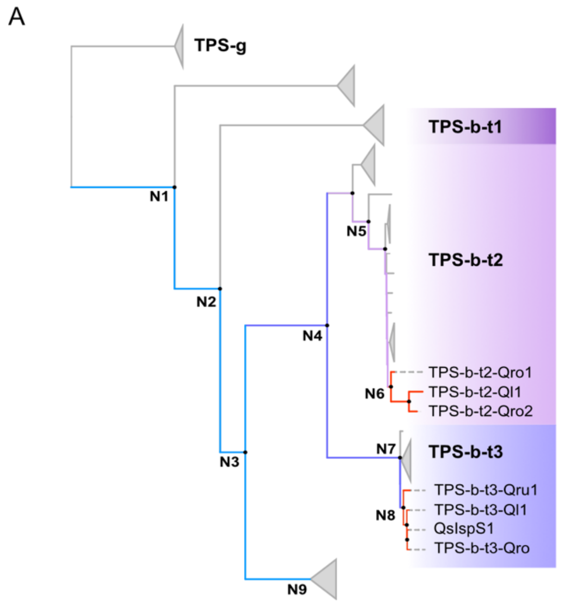


B

| Node | F338 | S/V446 | F485 | N505 | Isoprene score |
|---|---|---|---|---|---|
| N1 | I | V | L | N | 2 |
| N2 | I | V | L | N | 2 |
| N3 | T | V | L | N | 2 |
| N4 | T | V | F | N | 3 |
| N5 | T | V | F | N | 3 |
| N6 | F | V | F | N | 4 |
| N7 | T | V | F | N | 3 |
| N8 | F | V | F | N | 4 |
| N9 | N | V | L | K | 1 |

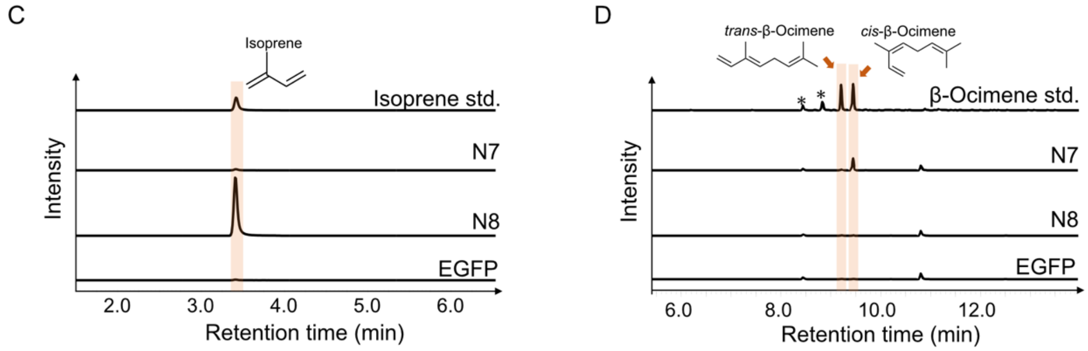


**Figure S7.** Results of ancestral sequence reconstruction of Fagaceae TPS genes using GRaSP. (*A*) Maximum-likelihood phylogeny of the TPS gene set used for ancestral reconstructions. The phylogeny was inferred with RAxML v8.2.11 under the GAMMA GTR substitution model, using 1,000 bootstrap replicates to assess node support. Internal nodes N1–N8 mark the bifurcation points analysed. Branch colours denote the inferred isoprene score—blue (2), purple (3) and red (4). The three focal clades (TPS-b-t1, TPS-b-t2 and TPS-b-t3) are highlighted in distinct colours for ease of reference. (*B*) Inferred amino-acid states at the diagnostic tetrad and the corresponding isoprene score for each ancestral node (N1–N8). Key amino acid changes were as follows: N4 – F485: Leu → Phe. N6 – F338: Thr → Phe. N8 – F338: Thr → Phe. (*C*) and (*D*) Enzyme assays of the ancestral sequence products using DMAPP and GPP as substrates. The reaction products were detected at *m/z* 67 for isoprene (*C*) and *m/z* 93 for monoterpenes (*D*) as the substrate. The *cis* or *trans* configurations of β-ocimene were identified based on spectral annotation. The chromatograms, except for the β-ocimene standard, are plotted on the same scale. Asterisks represent peaks of impurities.

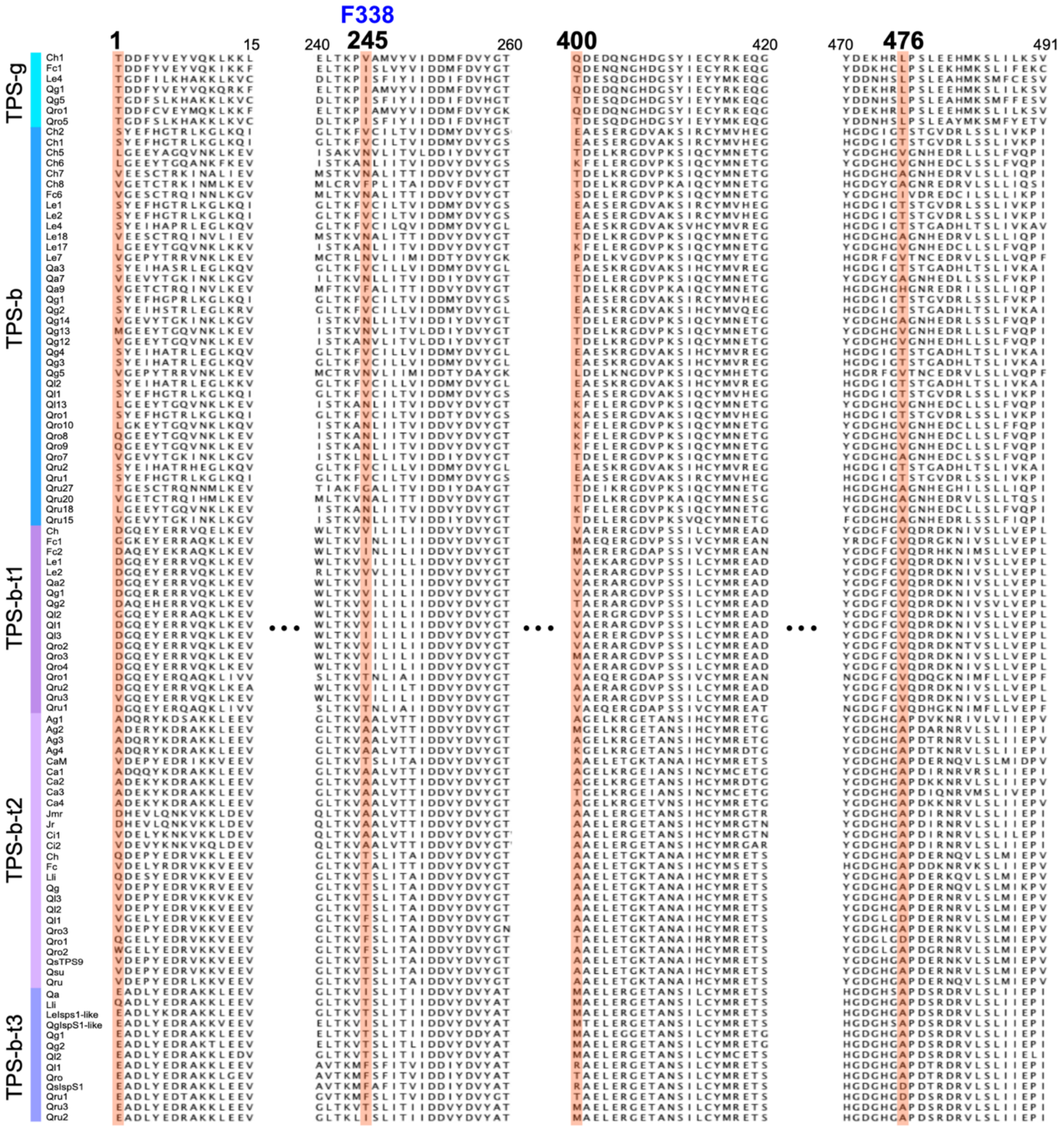

**Figure S8.** Multiple sequence alignment of TPS-b and TPS-g subfamily members from 17 Fagales species. Four candidate sites under positive selection are highlighted by red. The 245th site corresponds to F338, one conservative amino acid residue for isoprene synthases.

**Table S1.** Summary statistics for the genomes of the eight analyzed Fagaceae species.

| Species | Number of chromosomes | Genome size (Mb) | Gene count | Reference |
|---|---|---|---|---|
| *Q. robur* | 12 | 789.2 | 45,167* | NCBI, assembly dhQueRobu3.1 (Bodénès et al., 2016) |
| *Q. lobata* | 12 | 845.9 | 45,898* | NCBI, assembly ValleyOak3.2 (Sork et al., 2016) |
| *Q. rubra* | 12 | 739 | 33,247* | NCBI, assembly Qrubra_687_v2.0 (Kapoor et al., 2023) |
| *Q. acutissima* | 12 | 758.1 | 31,040 | Genome Warehouse (Fu *et al.*, 2022) |
| *Q. glauca* | 12 | 893.5 | 39,758 | Kudo et al., 2025 |
| *L. edulis* | 12 | 861.6 | 34,059 | Kudo et al., 2025 |
| *C. hystrix* | 12 | 882.7 | 37,750 | Huang et al., 2023 |
| *F. crenata* | 12 | 539.1 | 35,116 | Torimaru et al., 2025 |

Asterisks indicate that gene counts include isoforms.

**Table S2.** Isoprene emission factors reported in previous studies.

| Species | Isoprene emission factor | Reference |
|---|---|---|
| *Q. robur* | 79.3 (μgg-1h-1) | Keenan et al., 2009 |
| *Q. lobata* | 86 (μgCg-1h-1) | Geron et al., 2001 |
| *Q. rubra* | 58.2 (μgg-1h-1) | Keenan et al., 2009 |
| *Q. acutissima* | 0.038 - 0.078 (μgCgdw-1h-1) | Lim et al., 2011 |
| *Q. glauca* | 0.04 (μggdw-1h-1) | Bao et al., 2008 |
| *Q. serrata* | 224.21 (μggdw-1h-1) | Bao et al., 2008 |
| *L. edulis* | No emit | Mochizuki et al., 2020 |
| *C. hystrix* | 1.91 ± 0.10 (μgg−1h−1) | Zeng et al., 2022 |
| *F. crenata* | 0.79 (μggdw-1h-1) | Bao et al., 2008 |

'dw' indicates dry weight.

**Table S3**. List of the sequences used for ancestral amino acid sequence inference and detection of positively selected residues.

| Gene ID | Accession identified by blast search against database (Fagales) | Group |
|---|---|---|
| TPSb-t1, t2, t3 (Isps candidate clades) 57 sequences | | |
| TPS-b-t1-Ch | | TPS-b-t1 |
| TPS-b-t1-Fc1 | | TPS-b-t1 |
| TPS-b-t1-Fc2 | | TPS-b-t1 |
| TPS-b-t1-Le1 | | TPS-b-t1 |
| TPS-b-t1-Le2 | | TPS-b-t1 |
| TPS-b-t1-Qa2 | | TPS-b-t1 |
| TPS-b-t1-Qg1 | | TPS-b-t1 |
| TPS-b-t1-Qg2 | | TPS-b-t1 |
| TPS-b-t1-Ql2 | | TPS-b-t1 |
| TPS-b-t1-Ql1 | | TPS-b-t1 |
| TPS-b-t1-Ql3 | | TPS-b-t1 |
| TPS-b-t1-Qro2 | | TPS-b-t1 |
| TPS-b-t1-Qro3 | | TPS-b-t1 |
| TPS-b-t1-Qro4 | | TPS-b-t1 |
| TPS-b-t1-Qro1 | | TPS-b-t1 |
| TPS-b-t1-Qru2 | | TPS-b-t1 |
| TPS-b-t1-Qru3 | | TPS-b-t1 |
| TPS-b-t1-Qru1 | | TPS-b-t1 |
| TPS-b-t2-Jr | XP_035539955 | TPS-b-t2 |
| TPS-b-t2-Jmr | XM_041148044 | TPS-b-t2 |
| TPS-b-t2-Ag1 | XP_062151842 | TPS-b-t2 |

| | | |
|---|---|---|
| TPS-b-t2-Ca1 | XP_059450886 | TPS-b-t2 |
| TPS-b-t2-Ca2 | XP_059451578 | TPS-b-t2 |
| TPS-b-t2-Ca3 | XP_059451982 | TPS-b-t2 |
| TPS-b-t2-Ca4 | XP_059452022 | TPS-b-t2 |
| TPS-b-t2-Ci2 | XP_042971821 | TPS-b-t2 |
| TPS-b-t2-Ci1 | XP_042971785 | TPS-b-t2 |
| TPS-b-t2-Ag2 | XP_062153083 | TPS-b-t2 |
| TPS-b-t2-Ag3 | XP_062153084 | TPS-b-t2 |
| TPS-b-t2-Ag4 | XP_062153149 | TPS-b-t2 |
| TPS-b-t2-Fc | | TPS-b-t2 |
| TPS-b-t2-Ch | | TPS-b-t2 |
| TPS-b-t2-CaM | | TPS-b-t2 |
| TPS-b-t2-Lli | KAL0001771 | TPS-b-t2 |
| TPS-b-t2-Qg | | TPS-b-t2 |
| TPS-b-t2-Ql3 | | TPS-b-t2 |
| TPS-b-t2-Ql2 | | TPS-b-t2 |
| TPS-b-t2-Ql1 | | TPS-b-t2 |
| TPS-b-t2-Qro3 | | TPS-b-t2 |
| TPS-b-t2-Qro1 | | TPS-b-t2 |
| TPS-b-t2-Qro2 | | TPS-b-t2 |
| QsTPS_score2 | | TPS-b-t2 |
| TPS-b-t2-Qsu | XP_065622397 | TPS-b-t2 |
| TPS-b-t2-Qru | | TPS-b-t2 |
| TPS-b-t3-Lli | KAL0001769 | TPS-b-t3 |
| Lthed_cloned_aa | | TPS-b-t3 |
| TPS-b-t3-Qa | | TPS-b-t3 |

| | | |
|---|---|---|
| Qgl_cloned_aa | | TPS-b-t3 |
| TPS-b-t3-Qg1 | | TPS-b-t3 |
| TPS-bt-t3-Qg2 | | TPS-b-t3 |
| TPS-b-t3-Ql2 | | TPS-b-t3 |
| TPS-b-t3-Ql1 | | TPS-b-t3 |
| TPS-b-t3-Qro | | TPS-b-t3 |
| QsIspS1 | | TPS-b-t3 |
| TPS-b-t3-Qru1 | | TPS-b-t3 |
| TPS-b-t3-Qru3 | | TPS-b-t3 |
| TPS-b-t3-Qru2 | | TPS-b-t3 |
| TPSb others 38 sequences | | |
| TPS-b-Ch5 | | TPS-b |
| TPS-b-Ch6 | | TPS-b |
| TPS-b-Ch7 | | TPS-b |
| TPS-b-Ch8 | | TPS-b |
| TPS-b-Fc6 | | TPS-b |
| TPS-b-Le18 | | TPS-b |
| TPS-b-Le17 | | TPS-b |
| TPS-b-Qa7 | | TPS-b |
| TPS-b-Qa9 | | TPS-b |
| TPS-b-Qg14 | | TPS-b |
| TPS-b-Qg13 | | TPS-b |
| TPS-b-Qg12 | | TPS-b |
| TPS-b-Qg5 | | TPS-b |
| TPS-b-Le7 | | TPS-b |
| TPS-b-Ql13 | | TPS-b |

| | | |
|---|---|---|
| TPS-b-Qro10 | | TPS-b |
| TPS-b-Qro8 | | TPS-b |
| TPS-b-Qro9 | | TPS-b |
| TPS-b-Qro7 | | TPS-b |
| TPS-b-Qru27 | | TPS-b |
| TPS-b-Qru20 | | TPS-b |
| TPS-b-Qru18 | | TPS-b |
| TPS-b-Qru15 | | TPS-b |
| TPS-b-Ch2 | | TPS-b |
| TPS-b-Ch1 | | TPS-b |
| TPS-b-Le1 | | TPS-b |
| TPS-b-Le2 | | TPS-b |
| TPS-b-Le4 | | TPS-b |
| TPS-b-Qa3 | | TPS-b |
| TPS-b-Qg1 | | TPS-b |
| TPS-b-Qg2 | | TPS-b |
| TPS-b-Qg4 | | TPS-b |
| TPS-b-Qg3 | | TPS-b |
| TPS-b-Ql2 | | TPS-b |
| TPS-b-Ql1 | | TPS-b |
| TPS-b-Qro1 | | TPS-b |
| TPS-b-Qru2 | | TPS-b |
| TPS-b-Qru1 | | TPS-b |
| TPSg 7 sequences | | |
| TPS-g-Ch1 | | TPS-g |
| TPS-g-Fc1 | | TPS-g |

| | | |
|---|---|---|
| TPS-g-Qg1 | | TPS-g |
| TPS-g-Qro1 | | TPS-g |
| TPS-g-Qg5 | | TPS-g |
| TPS-g-Le4 | | TPS-g |
| TPS-g-Qro5 | | TPS-g |

Qro; *Quercus robur* (Fagaceae) gain from genomic survey.
Qg; *Quercus glauca* (Fagaceae) gain from genomic survey.
Qru; *Quercus rubra* (Fagaceae) gain from genomic survey.
Qa; *Quercus acutissima* (Fagaceae) gain from genomic survey.
Qs; *Quercus serrata* (Fagaceae) gain from genomic survey.
Le; *Lithocarpus edulis* (Fagaceae) gain from genomic survey.
Ch; *Castanopsis hystrix* (Fagaceae) gain from genomic survey.
Fc; *Fagus crenata* (Fagaceae) gain from genomic survey.
Ql; *Quercus lobata* (Fagaceae) gain from genomic survey.
Qsu; *Quercus suber* (Fagaceae) gain from BlastP search against database.
CaM; *Castanea mollissima* (Fagaceae) gain from BlastP search against database.
Lli; *Lithocarpus litselfolius* (Fagaceae) gain from BlastP search against database.
Jr; *Juglans regia* (Juglandaceae) gain from BlastP search against database.
Jmr; *Juglans maicrocarpa x Juglans regida* (Juglandaceae) gain from BlastP search against database.
Ci; *Carya illinoinensis* (Juglandaceae) gain from BlastP search against database.
Ca; *Corylus avellana* (Betulaceae) gain from BlastP search against database.
Ag; *Alnus glutinosa* (Betulaceae) gain from BlastP search against database.
Lthed_cloned_aa; *Lithocarpus edulis* (Fagaceae) identified by cloning.
Qgl_cloned_aa; *Quercus glauca* (Fagaceae) identified by cloning.

**Table S4.** Candidate sites under positive selection identified under branch-site model.

| Model | Np | Log-likelihood | Estimates of parameters | | | | | LRT $P$-value | Positive sites, BEB probability |
|---|---|---|---|---|---|---|---|---|---|
| Model A | 207 | -31524.908 | Site class | 0 | 1 | 2a | 2b | $1.027\times10^{-6}$ | 1, 0.964<br>245, 0.953<br>400, 0.966<br>476, 0.993 |
| | | | Frequency | 0.700 | 0.263 | 0.027 | 0.010 | | |
| | | | Background ω | $\omega_0$= 0.252 | $\omega_1$= 1.000 | $\omega_0$= 0.252 | $\omega_1$= 1.000 | | |
| | | | Foreground ω | $\omega_0$= 0.252 | $\omega_1$= 1.000 | $\omega_2$= 8.912 | $\omega_2$= 8.912 | | |
| Model A null | 206 | -31536.846 | | | | | | | Not Allowed |

The clades that acquired F338 were predefined as foreground branches.

Np; number of free parameters

Site class 0; under purifying selection along all branches with $0 < \omega_0 < 1$

Site class 1; undergoing neutral evolution along all branch with $\omega_1 = 1$

Site class 2a; positive selection along foreground branches with $\omega_2 > 1$, while the background branches are under purifying selection with $0 < \omega_0 < 1$

Site class 2b; positive selection along foreground branches with $\omega_2 > 1$, while the background branches are undergoing neutral evolution with $\omega_1 = 1$

Positive sites; Sites that have a probability higher than 95% to be under positive selection according to the Bayes Empirical Bayes (BEB) method. The numbers refer to site positions in the gap-free alignment used for the test. Site position 245 correspond to F338.

**Table S5.** Primers used in this study.

| Name | Sequence (5’- 3’) |
|---|---|
| N7_Fw | AAGGAGATATACATAATGGCAAGCAAACAGGTTCTG |
| N7_grasp_Rv | GCTCGAATTCGGATCCAGATGAATATGGCTGTGAAAATTG |
| N7_paml_Rv | GCTCGAATTCGGATCCAGATGGATCTGACTATGAAAATTG |
| N8_paml_Fw | ACCAGCATGGCCGAACTGGAACGTGGTG |
| N8_paml_Rv | TTCGGCCATGCTGGTGCCCAGATCATTACAC |
| QrIspS_Fw | AAGGAGATATACATAATGGCAAGCGCACCGACC |
| QrIspS_Rv | GCTCGAATTCGGATCAATCAGGCTCACCGGTTCAATC |

**Dataset S1**

Multiple sequence alignment of ancestral sequences of Fagaceae TPS genes using PAML and GRaSP. The four conserved amino acid residues used for calculating isoprene score were shown by black frame.
