## Supplementary figures and images for "Molecular evolution of terpene synthase underlying the diversification of isoprene emission in Fagaceae"

### Dataset S1

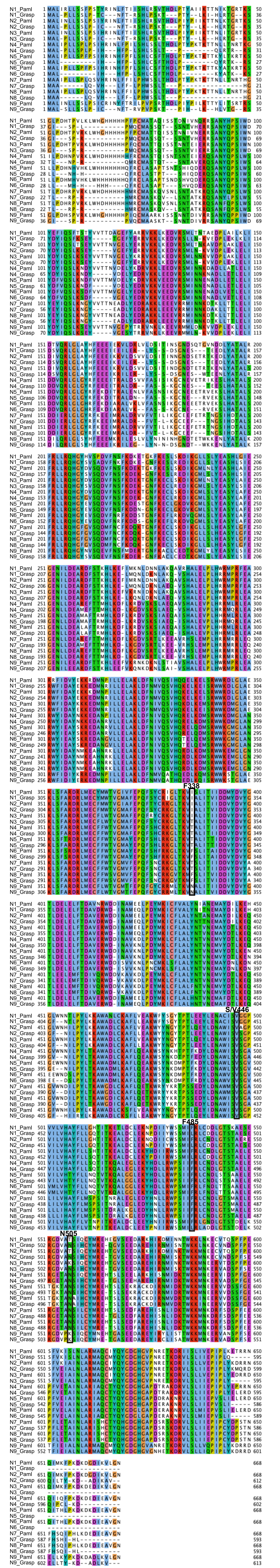
